# Network Modeling Predicts How DYRK1A Inhibition Promotes Cardiomyocyte Cycling after Ischemic/Reperfusion Injury

**DOI:** 10.1101/2025.08.19.671147

**Authors:** Bryce C. Murillo, Alexander Young, Kaitlyn L. Wintruba, Alexander J. Eichert, Klara Siejda, Dennon Hoenig, Leigh A. Bradley, Bryana N. Harris, Catherine Zhao, Michelle Wu, Emmanuel Deau, Matthias F. Lindberg, Laurent Meijer, Jeffrey J. Saucerman, Matthew J. Wolf

## Abstract

The adult mammalian heart has a limited ability to regenerate lost myocardium following myocardial infarction (MI), largely due to the poor proliferative capacity of cardiomyocytes (CMs). Dual-specificity tyrosine phosphorylation-regulated kinase 1A (DYRK1A) is a known regulator of cell quiescence, though the mechanisms underlying its function remain unclear. Previous studies have shown that pharmacological inhibition of DYRK1A using harmine induces CM cell cycle re-entry after ischemia/reperfusion (I/R) MI. Here, we developed a computational network model of DYRK1A-mediated regulation of the cell cycle, which predicts how DYRK1A inhibition promotes CM re-entry. To validate these predictions, we tested selective DYRK1A inhibitors and observed robust induction of cell cycle activity in neonatal rat cardiomyocytes (NRCMs). Integrating our network model with bulk RNA-sequencing data from DYRK1A inhibitor-treated NRCMs, we identified E2F1 as a key transcriptional driver of cell cycle gene expression. Finally, we demonstrate that both pharmacological and post-developmental inhibition of DYRK1A enhances heart function and increases CM cycling following I/R MI. Our findings suggest that functional recovery induced by small molecule inhibitor of DYRK1A is mediated by the induction of cycling CMs.

**One Sentence Summary:** Inhibition of DYRK1A through LCTB-92 induces cardiomyocyte cycling and improved heart function in a mouse model of ischemic/reperfusion injury.

## INTRODUCTION

Myocardial infarctions (MIs) affect ∼800,000 individuals in the United States annually and cause significant morbidity and mortality (*1*). The ischemia induced by the cessation of coronary blood flow during an MI and subsequent reperfusion leads to cardiomyocyte (CM) necrosis, infiltration of inflammatory cells, and activation of fibroblasts to produce fibrosis (*2*). Unfortunately, the adult mammalian heart has no significant regenerative capacity to restore CM loss after MI (*3, 4*). Therefore, the current limitations in treating MIs and the need for new therapeutics to stimulate CM proliferation, in a controlled manner, represent an opportunity to improve cardiac function after injury.

A number of studies demonstrate that enhancing CM proliferation after MI improves heart function (*5–7*). The genetic ablation or pharmacologic inhibition of proteins in the YAP (Yes-associated protein) pathway promotes CM cycling. For example, the genetic ablation of the Salvador scaffold protein or MST1 and MST2, two serine/threonine kinases that regulate YAP, promote robust CM proliferation (*8, 9*). Alternatively, the overexpression of cyclin-dependent kinase 1 (CDK1), CDK4, cyclin B1, and cyclin D1 or CDK4 and cyclin D1 in combination with Wee1 and TGFβ inhibition promotes CM proliferation in vivo (*10*). We previously identified that genetic ablation or inhibition of dual specificity tyrosine phosphorylation regulated kinase 1A (DYRK1A) induced CM cell cycle entry and improved heart function after ischemic reperfusion (I/R) MI (*11*). DYRK1A is an upstream mediator of the Myb-MuvB/dimerization partner, RB-like, E2F, and multi-vulval class B (DREAM) complex, which maintains cell cycle quiescence through suppression of G1/S and G2/M checkpoints (*12*). DYRK1A promotes DREAM complex assembly through phosphorylation of LIN52 on residue serine 28 (*12*). DYRK1A also phosphorylates cyclin D2, targeting it for proteasomal degradation in CMs (*13*). In contrast, the overexpression of DYRK1A decreased CM proliferation during development in a mouse model of Down Syndrome (*14*). This provides evidence that DYRK1A regulates cell cycle quiescence, and its inhibition can promote re-entry of CMs into the cell cycle. However, the mechanisms through which DYRK1A influences cell cycling remain enigmatic.

Our prior experiments have utilized Harmine, an inhibitor of DYRK1A that also inhibits Monoamine Oxidase (MAO) (*11*). While Harmine has proven effective in promoting CM cycling, it is not without its drawbacks. The drug’s adverse effects, including nausea, vomiting, drowsiness, and impaired concentration, significantly limit its potential clinical use (*15*). Therefore, our current efforts are focused on developing more specific, orally bioavailable DYRK1A inhibitors that can effectively promote CM cycling without these adverse effects, thereby paving the way for their potential clinical application.

Here, we employed computational network modeling to predict mechanisms by which DYRK1A regulates CM cell cycling. These predictions were validated in neonatal rat cardiomyocytes (NRCMs) using selective DYRK1A inhibitors, followed by RNA-sequencing. Based on these findings, we tested the DYRK1A inhibitor Leucettinib-92 (LCTB-92) in a mouse model of ischemia-reperfusion (I/R) myocardial infarction (MI), where it improved heart function and increased CM cycling.

## RESULTS

### Network Model Predicts Mechanisms of DYRK1A-mediated Cell Cycle Activity

While we have previously found that both genetic and pharmacologic inhibition of DYRK1A induce CM cycling (*11*), the mechanisms by which they do so have not been explored. This led us to develop a network model by manual curation of literature that describes DYRK1A’s regulatory role of cell cycling molecules (**Table S1**). DYRK1A’s role in cell cycling is believed to be highly conserved across different tissue types, so we broadened our literature review beyond papers that focused on cardiomyocytes to leverage mechanistic knowledge from other cell types (*12, 16–21*). We then used these relationships to create a logic-based network model using Netflux (*22, 23*), where we could mechanistically predict how DYRK1A knockdown increases cell cycle activity (**Table S2**).

The model predicts that in quiescent CMs (baseline), high DYRK1A expression induces DREAM complex formation and RB-1 activity through the direct and indirect degradation of cyclin D and E respectively (**Fig. 1A**). Upon simulated DYRK1A knockdown, the model predicts an increase in cell cycle entry (DNA replication) through the increase of cyclins D and E, which promotes DREAM complex breakdown and inactivation of RB-1 (**Fig. 1A**). We then validated our model’s accuracy by comparing predictions with experimental results found in literature not used to build the model (**Fig. 1B & C, Table S3**). Overall, the model’s predictions were highly accurate across multiple experiments performed in cardiomyocyte (100%) and cancer (85%) literature (*11, 13, 24–40*). Interestingly, the model’s predicted effect of DYRK1A KO on DNA replication was consistent with cardiomyocyte literature but not consistent with cancer literature (*31, 38*). DYRK1A may play an alternative role in the context of some cancers (*41*) which may explain this discrepancy in literature.

**Figure 1:**
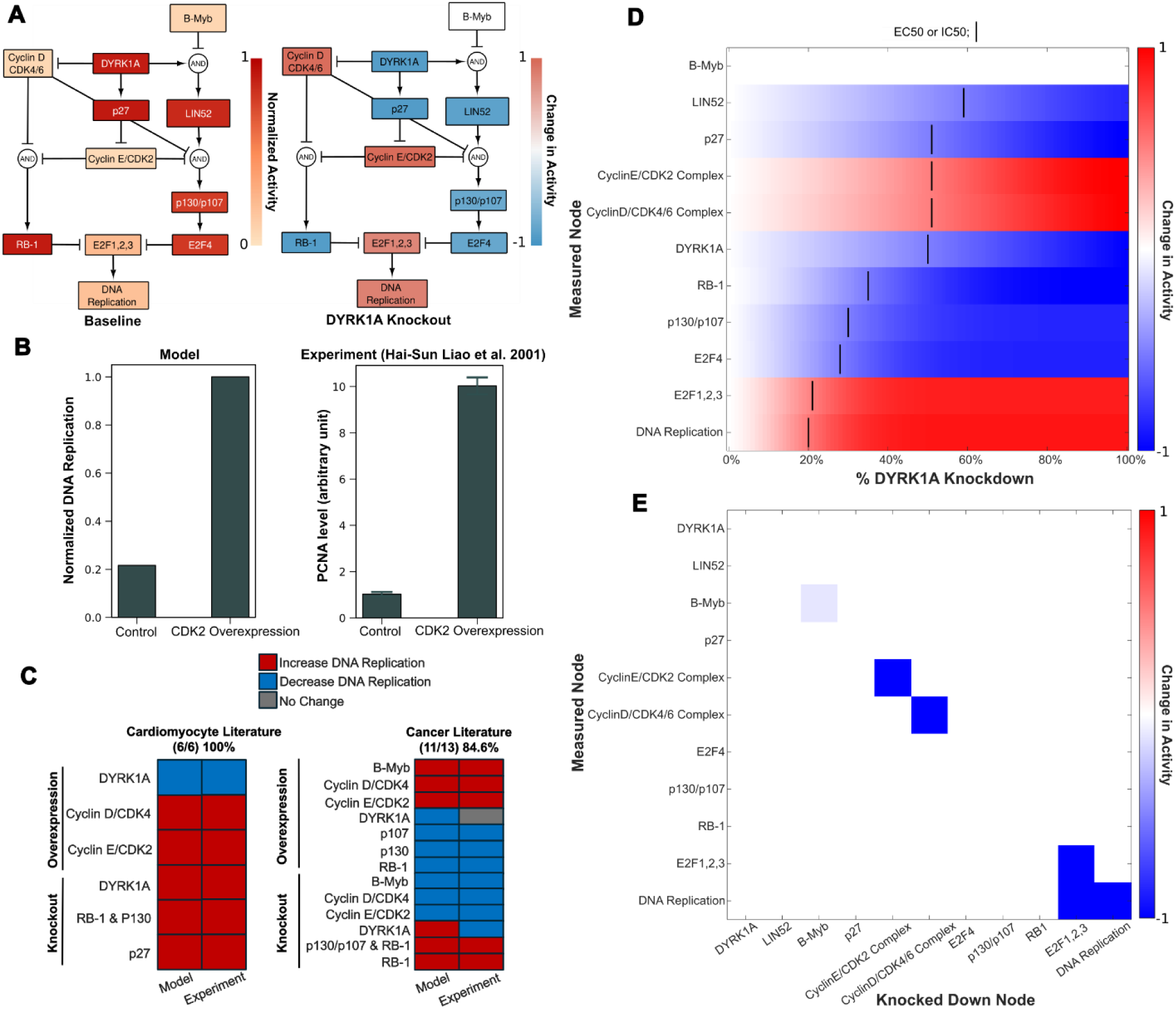
Network model predicts mechanisms of DYRK1A-mediated cardiomyocyte cell cycle activity. **(A)** Model predicts response to active DYRK1A (left) and when DYRK1A is knocked out (right). **(B)** Model simulation of CDK2 overexpression’s effect on DNA replication (left) compared to experimental data found in literature (right). **(C)** Other simulations were compared to experiments in both cardiomyocyte (n = 5) and cancer literature (n = 13) (n = # of primary literature articles). Validation accuracy is measured by the # of predictions that match literature results/simulations performed. **(D)** Concentration response showing resulting node activity from increasing levels of DYRK1A knockdown. IC50 and EC50 values indicate the % DYRK1A knockdown at which a node increases or decreases by 50% (determined by the change in normalized activity by 50%). **(E)** Knockdown screen showing network model’s response to each node being 100% knocked down. Nodes were knocked out one at a time by setting their y-max to 0. Simulations were performed under a 100% DYRK1A knockdown.

To identify which network nodes are most sensitive to DYRK1A, we simulated a DYRK1A expression concentration response. Node sensitivity was assessed based on predicted EC50/IC50 values (**Fig. 1D**). Strikingly, nodes further downstream targets were more sensitive to DYRK1A knockdown, indicating that even partial DYRK1A inhibition may be sufficient to induce cell cycle entry. Finally, we performed a knockdown screen in the context of a 100% DYRK1A knockdown to identify modulators of its response (**Fig. 1E**). We found that knockdown of E2F1,2, and 3 transcription factors attenuated the predicted increase in DNA replication upon a DYRK1A knockdown. Additionally, knockdown of cyclin D and E complexes did not attenuate the effect of DYRK1A knockdown on DNA replication, indicating that they may play redundant roles in modulating cell cycle entry. All in all, our robust analysis supports that we have built a reliable model of DYRK1A signaling.

### DYRK1A inhibition promotes cell cycling in neonatal rat cardiomyocytes

To experimentally validate the network model prediction that DYRK1A inhibition upregulates cyclin D and promotes cell cycle entry, we tested recently reported DYRK1A inhibitors, LCTB-21 and LCTB-92, in NRCMs (*42, 43*). LCTB-92 and LCTB-21 work by binding to the active site of DYRK1A and preventing kinase activity. Inactive forms of both compounds (ISO-92 and ISO-21, respectively) were used as negative controls along with DMSO. High-content imaging and segmentation of α-actinin, Ki67, cyclin D1, and DAPI (**Fig. 2A**) were used to analyze CM cell cycle activity at single-cell resolution. The network model predicted that DYRK1A inhibition should increase DNA replication in a dose-dependent manner (**Fig. 2B**). Consistent with these predictions, treatment with higher concentrations of LCTB-21 (1 µM) and LCTB-92 (0.1 and 1 µM) increased cell cycle activity as evidenced by a higher percentage of Ki67-positive CMs compared to DMSO and isomeric controls (**Fig. 2C**). The model also predicted a dose-dependent increase in cyclin D levels following DYRK1A inhibition (**Fig. 2D**). Supporting the model predictions, validation experiments showed an increase in cyclin D1 expression in CMs treated with LCTB-21 (1 µM) and LCTB-92 (0.1 and 1 µM) compared to DMSO and isomeric controls (**Fig. 2E**).

**Figure 2:**
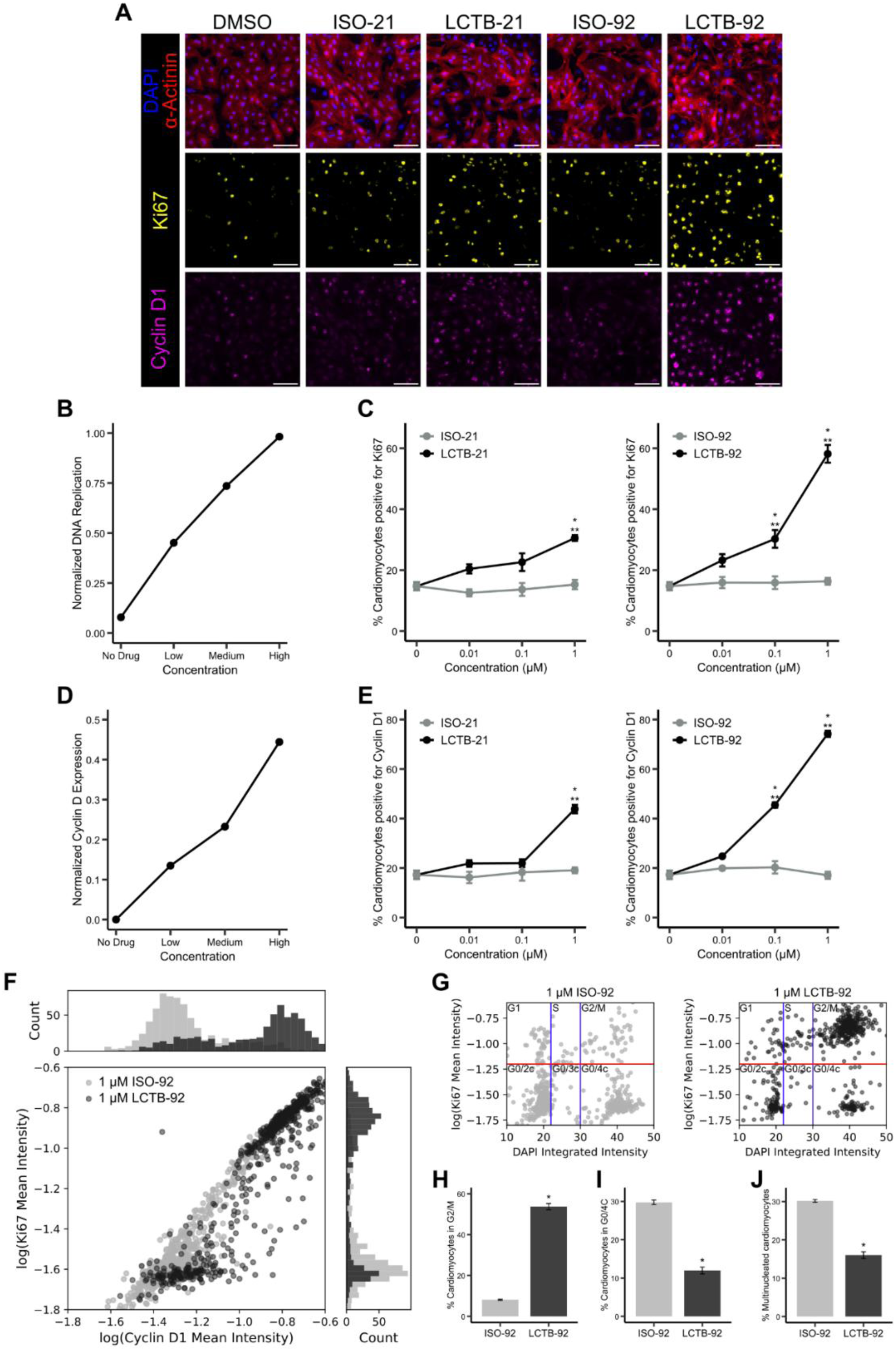
DYRK1A inhibitor LCTB-92 increases cell cycle activity of neonatal rat cardiomyocytes, validating network model. **A)** Representative images of cardiomyocytes treated with DYRK1A inhibitors (LCTB-21, LCTB-92) or controls (DMSO, ISO-21, ISO-92) and labeled with DAPI (blue), α-actinin (red), Ki67 (yellow), and cyclin D1 (magenta). Scale bar = 100 mm. **B)** Network model prediction of DNA replication and **C)** experimental quantification of the percent of cardiomyocytes expressing Ki67 in response to DYRK1A inhibition. **D)** Network model prediction of cyclin D expression and **E)** experimental quantification of the percentage of cardiomyocytes expressing cyclin D1 in response to DYRK1A inhibition. **F)** Single-cell correlation between Ki67 and cyclin D1 expression levels with histograms illustrating the distribution of intensities for each marker. **G)** Bivariate analysis of DNA content and Ki67 to separate out cell cycle phase. DNA content thresholds separating 2c, 3c, and 4c nuclei are shown as blue vertical lines. Ki67 positive threshold is shown as a red horizontal line. Percent of cardiomyocytes in **H)** G2/M and **I)** G0/4c phases from panel G. **J)** Percent of multinucleated cardiomyocytes. Error bars represent mean ± SEM. Statistical significance for C and E was determined by one-way ANOVA followed by Tukey’s multiple comparisons test. \*\**p* < 0.05 LCTB treatments compared to DMSO (0 mM) control; \**p* < 0.05 LCTB treatments compared to the respective isomeric control. Statistical significance for H-J was determined by t-test. \**p* < 0.05.

Given the substantial increase in both Ki67 and cyclin D1 expression in CMs treated with 1 µM LCTB-92, we further investigated the relationship between these markers and the cell cycle phase under this condition. Analysis of the mean Ki67 and cyclin D1 intensities revealed a shift from a unimodal distribution in cells treated with ISO-92 to a bimodal distribution in LCTB-92-treated CMs (**Fig. 2F**). The shift suggests a discrete bifurcation in Ki67 and cyclin D1 expression that creates distinct subpopulations of CMs (*44*). Analysis of DNA content and Ki67 (**Fig. 2G**) revealed an increased proportion of CMs in the G2/M phase (**Fig. 2H**) and a decreased proportion in the G0/4c (**Fig. 2I**) and G0/2c phases (*c* refers to DNA content, with 2c and 4c representing diploid and tetraploid levels, respectively) following LCTB-92 treatment compared to ISO-92. This shift toward late cell cycle progression, along with a reduction in multinucleation (**Fig. 2J**) suggests that LCTB-92 promotes true cell division rather than incomplete cell cycle progression.

To uncover the mechanistic differences that explain the efficacy of LCTB-92 vs. LCTB-21 in promoting CM proliferation, we performed RNA-sequencing on drug-treated NRCMs. Principal component analysis captured 88.6% of the variance across samples and revealed distinct clustering of our LCTB-92 and −21 treated groups (**Fig. 3A)**. Differential gene expression was performed in both drug-treated groups with respect to their inactive isomers (**Fig 3B-E).** Gene ontology (GO) enrichment analysis revealed that LCTB-92 treatment was associated with multiple terms related to cell cycle activity with “regulation of mitotic cell cycle” having the highest gene count (padj < 0.05) (**Fig 3B).** We found a much more pronounced effect in LCTB-92 treated cells with LCTB-92 having 1800 unique differentially expressed genes (DEGs), which could explain the more robust effect on cell cycling (**Fig 3E**).

**Figure 3:**
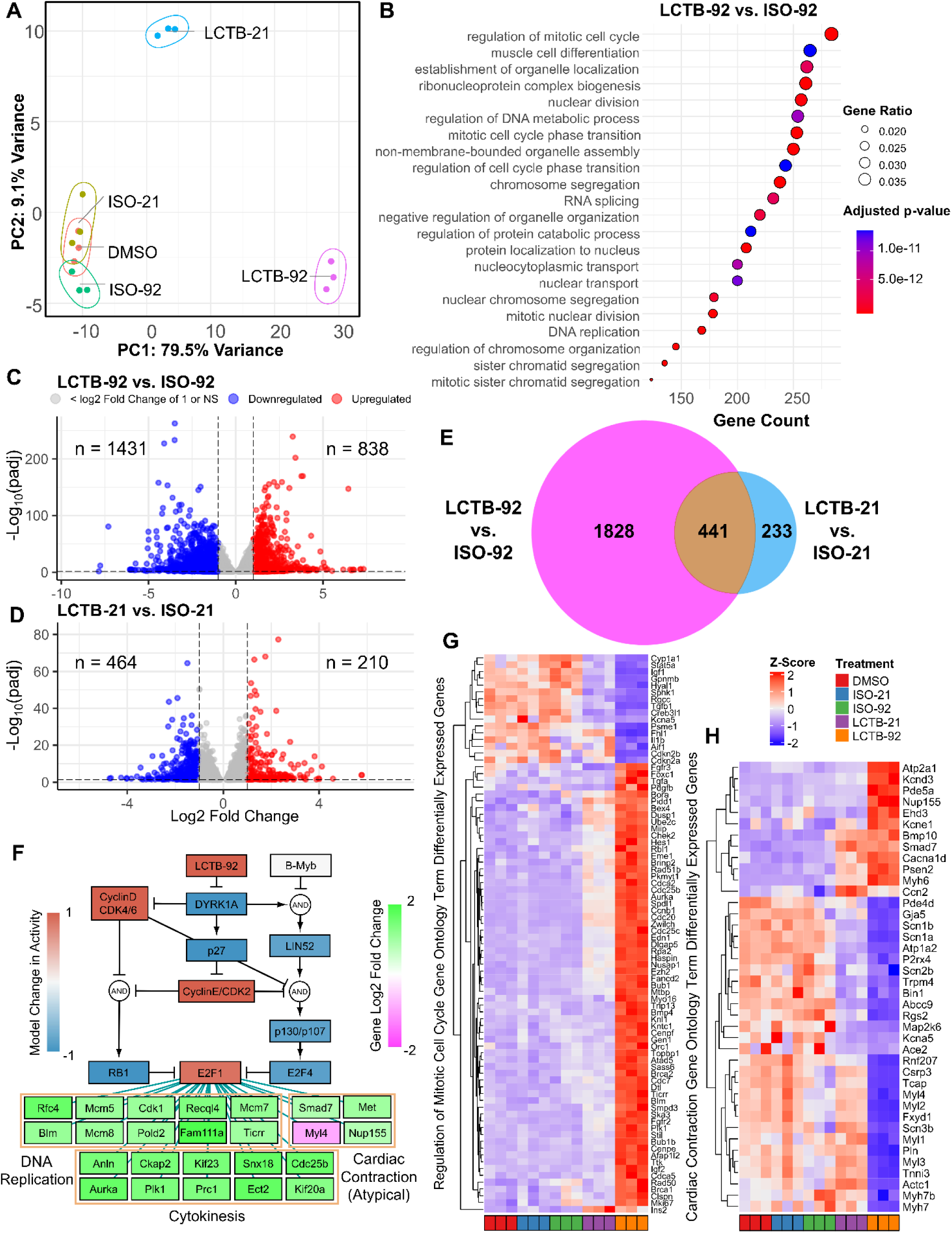
LCTB-92 treated NRCMs exhibit transcriptomic signatures associated with cell division. **(A)** Principal Component Analysis of drug-treated and respective control NRCM samples. **(B)** Gene Ontology Enrichment Analysis of LCT-92 treated NRCMs with respect to its inactive isomer (ISO-92). **(C-E)** Number of significantly differentiated expressed genes with ≥ 2-fold increase in LCTB-92 treated **(C)** or LCTB-21 treated **(D)** NRCMs compared to ISO-92 and ISO-21 respectively, and **(E)** the commonality between both treatment groups. **(F)** Network model simulation of LCTB-92 inhibition of DYRK1A and corresponding gene expression of cytokinesis, DNA replication, and cardiac contraction E2F1-target genes. **(G-H)** Z-score of differential gene expression in genes pulled from Cell Division **(G)** and Cardiac Contraction **(H)** Gene Ontology terms. Drug and isomeric controls groups were compared to DMSO for figures **G & H**. GO term enrichment analysis was performed with a p-value cutoff of 0.01 and q-value cutoff of 0.05.

To predict drivers of these changes in gene expression, we performed transcription factor enrichment using ChEA3 on the LCTB-92 DEGs (*45*). We found that E2F1 was a top hit according to mean library rank (**Table S4**). We then used the ChEA3 results to map the network model’s E2F1 node to DEGs associated with cell cycle related processes (**Fig. 3F**). Several E2F1 targets were also related to cardiac contraction, suggesting cardiomyocyte dedifferentiation which is a critical component of CM proliferation (*46*).

Despite the enrichment of its downstream targets, the normalized mRNA expression of E2F1, E2F2, and E2F3 were not significantly altered in LCTB-92 or LCTB-21–treated cells compared to their respective isomer controls (Fig. S1A–D). In contrast, analysis of ChEA3-predicted E2F1/2/3 target genes associated with mitotic processes revealed significant differential expression in both treatment groups compared to their respective isomer controls (Fig. S1E). Given that E2F activity is reported to be modulated at the protein level, our findings suggest that E2F transcription activity is modulated post-translationally in response to drug treatment (*18*). We then visualized the expression levels of genes involved in regulation of mitotic cell cycling and cardiac contraction, identified through our GO term enrichment analysis, across all treatment groups relative to DMSO (**Fig. 3G &H**). Notably, LCTB-92 treatment elicited a more pronounced transcriptional response, with increased expression of positive regulators and decreased expression of negative regulators of cell cycling. Additionally, we observed a greater decrease in cardiac contraction gene expression in the LCTB-92 treated samples, suggesting more robust cardiomyocyte dedifferentiation. Taken together, the integration of model predictions and transcriptomic data suggest LCTB-92 is inducing a more defined effect on cell cycling genes compared to LCTB-21, possibly through higher activation of E2F1.

### LCTB-92 promotes cardiomyocyte cell cycle activity and improves heart function post-MI

Based on the in vitro experimental results, we investigated the effects of LCTB-92 in an *in vivo* model of ischemic/reperfusion myocardial infarction (I/R MI). Specifically, we used our previously described αDKRC::RLTG transgenic mouse that restricts Cre recombinase expression to adult CMs that re-enter the cell cycle (*47*). The reporter mouse relies on the activation of the Ki67 promoter, providing an in vivo validation of the Ki67 results in our NRCM experiments. Cardiomyocytes that re-enter the cell cycle, based on the activation of a Ki67 promoter, express GFP. αDKRC::RLTG mice underwent I/R MI surgeries and were treated with 10 mg/kg LCTB-92 or ISO-92 via daily oral gavage for ten days (**Fig. 4A**). Serial echocardiography revealed that mice treated with LCTB-92 had significantly improved left ventricular ejection fractions (LVEFs) compared to ISO-92 treatment at both 2- and 4-weeks post-MI (**Fig 4B**). However, LCTB-92 treatment did not improve infarct size compared to control 4-weeks post-MI, despite improvement in heart function, suggesting the mechanism of improved cardiac function was not through infarct size reduction (**Fig 4. C & D).** Additionally, the hearts of mice treated with LCTB-92 had significantly increased number of cycled (GFP+) CMs compared to ISO-92 treatment (**Fig. 5A & B)**. Interestingly, LCTB-92 did not induce a significant increase of CM cycling in uninjured hearts, suggesting an injury-dependent context for the effects of LCTB-92 on CM cycling.

**Figure 4:**
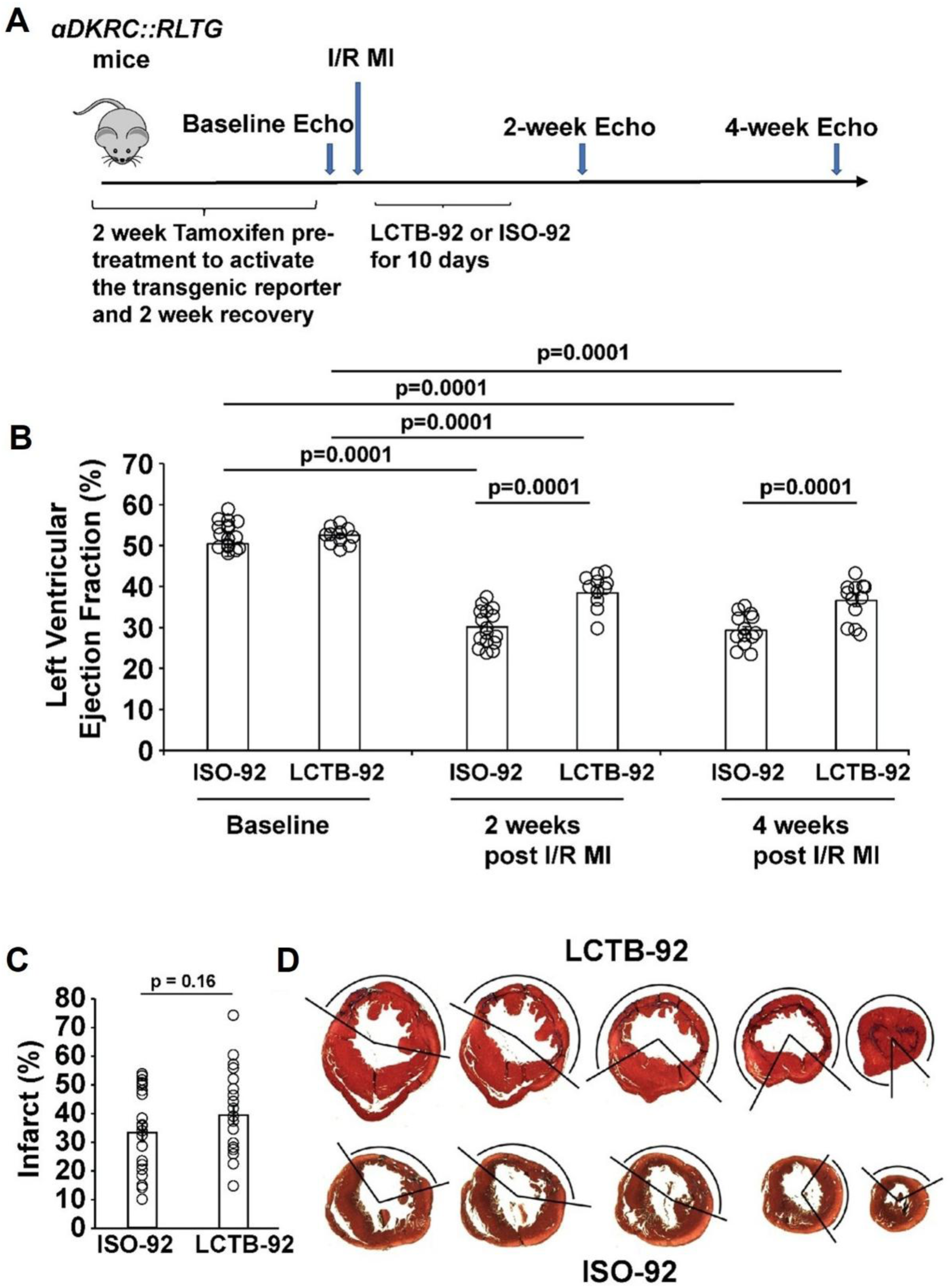
LCTB-92 improves heart function after I/R MI. **(A)** Schematic of 12-week-old αDKRC::RLTG mice that were treated with LCTB-92 or isomeric control for 10 days (10 mg/kg) following I/R MI. **(B)** Cardiac function measurements of αDKRC::RLTG mice treated with LCTB-92 or isomeric control after I/R MI, showing left ventricular ejection fraction (LVEF) 2- and 4-weeks post-MI. Error bars represent the mean ± SD and open circles are measurements for individual animals. N = 14 animals per group (9 males and 5 females). P-values were calculated using two-way ANOVA followed by Bonferroni’s multiple comparisons test. **(C)** Average infarct size (%) across whole heart sections from hearts treated with LCTB-92 or isomeric control 4 weeks after MI with **(D)** representative images. Error bars represent the mean ± SD and open circles are measurements for individual animals. Two samples t-test was performed for panel C.

**Figure 5:**
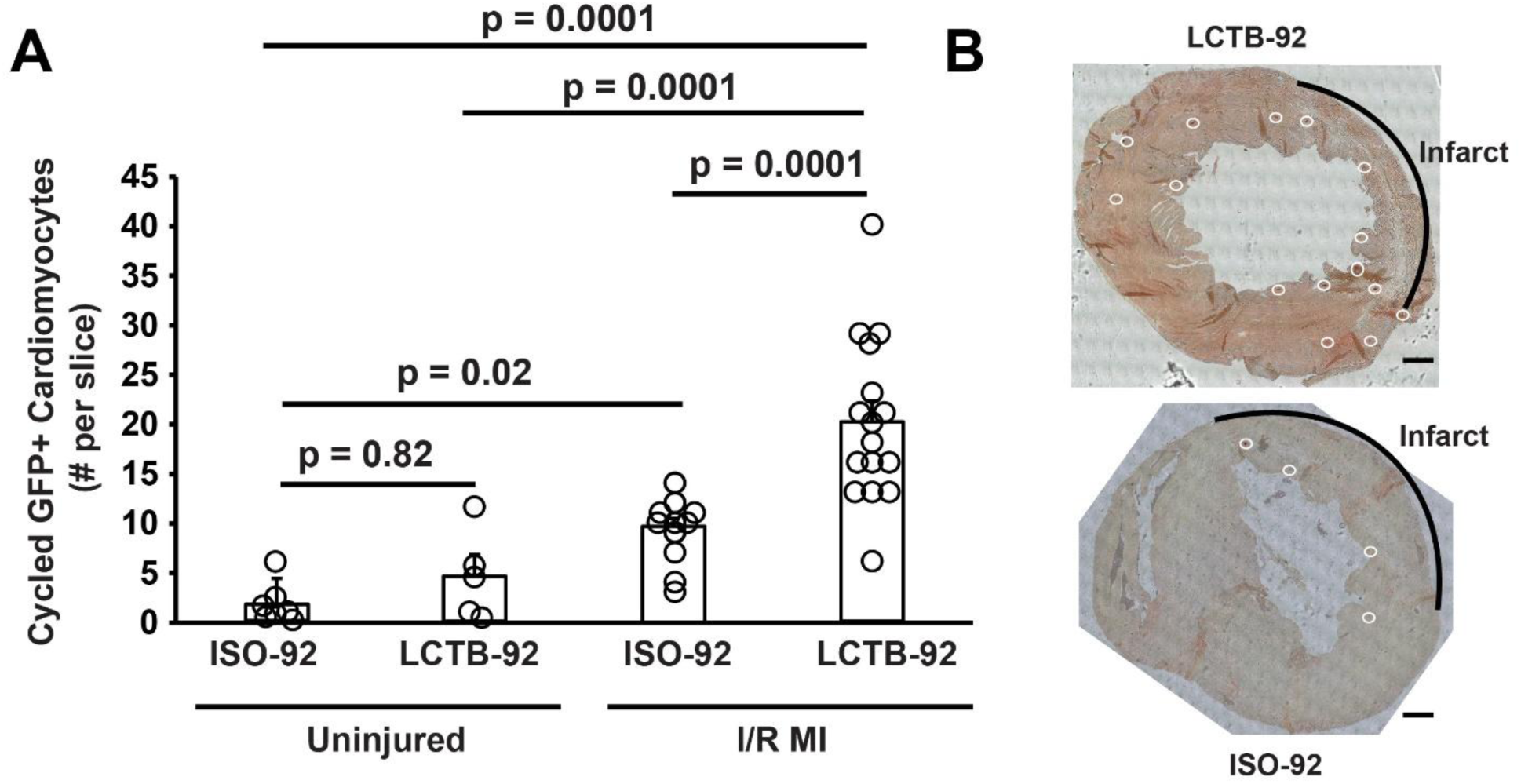
LCTB-92 promotes cardiomyocyte re-entry after I/R MI. (**A**) Quantification of the # of GFP^+^ cardiomyocytes collected from the midventricular section of hearts treated with LCTB-92 or isomeric control 4 weeks after MI with (**B**) Representative images of GFP^+^ cardiomyocytes indicating cell cycle re-entry. P-values were calculated using two-way ANOVA followed by Bonferroni’s multiple comparisons test. N = 7 animals for Iso-92 uninjured, N = 8 animals for LCTB-92 uninjured, N = 15 animals for ISO-92 I/R MI, and N = 16 animals for LCTB-92 I/R MI.

We then investigated if the improvement in heart function was linked to the increase in cell cycle activity induced by treatment with LCTB-92 using a previously described mouse model that ablates cardiomyocytes upon entry into the cell cycle through the Cre-mediated expression of Diphtheria toxin (*48*). αDKRC::RLTG/DTA or +::RLTG/DTA mice underwent I/R MI and were treated with 10 mg/kg LCTB-92 via daily oral gavage for ten days (**Fig. 6A**). Serial echocardiography showed that the effects of LCTB-92 were attenuated when cycling CMs were ablated after MI (**Fig 6. B-D**). Infarct sizes were similar between the two groups (**Fig 6. E & F**). Based on this, we would expect no change in ventricular function in uninjured mice treated with LCTB-92 due to there not being a significant increase in cycling CMs. Altogether, the beneficial effects of LCTB-92 on cardiac function after MI require increases in cycling CMs.

**Figure 6:**
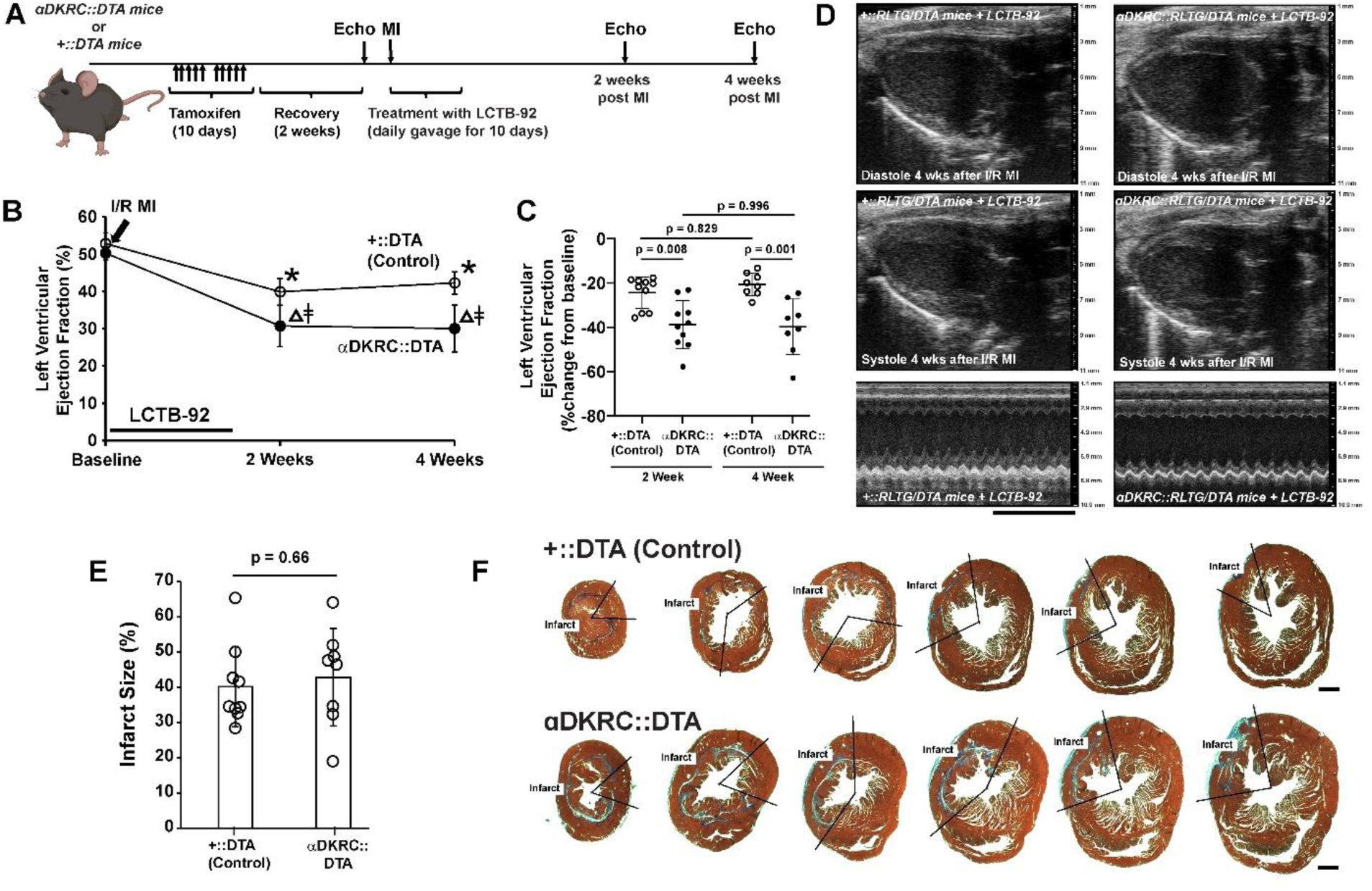
Ablation of cycling cardiomyocytes attenuates beneficial effects of LCTB-92 treatment. **(A)** Schematic of 12 week old αDKRC::RLTG/DTA or +::RLTG/DTA mice that were treated with LCTB-92 10 days (10 mg/kg) following I/R MI. **(B** and **C)** Cardiac function measurements of αDKRC::RLTG/DTA or control (+::RLTG/DTA) mice treated with LCTB-92 or isomeric control after I/R MI, showing absolute **(B)** absolute (%), and **(C)** percent change (% change from baseline) LVEF 2- and 4-weeks post-MI. N = 10 animal per group (5 male and 5 female) Values are mean +/− SD. p-values were calculated using two-way ANOVA followed by Bonferroni’s multiple comparisons test. * denotes p = 0.005 for each value compared to baseline. Δ denotes p=0.001 for each value compared to baseline. ǂ denotes p=0.05 for αDKRC::RLTG/DTA vs. control. **(D)** Representative echocardiography parasternal long-axis views of αDKRC::RLTG/DTA or control (+::RLTG/DTA) mice treated with LCTB-92 at 4 weeks after I/R MI. **(E)** Average infarct size (%) across whole heart sections from αDKRC::RLTG/DTA or control (+::RLTG/DTA) hearts treated with LCTB-92 at 4 weeks after MI with **(F)** representative images. Values are mean +/− SD. Two samples t-test was performed for panel E.

### The post-developmental genetic ablation of DYRK1A recapitulates beneficial effects of LCTB-92 treatment

Previously we observed aMHC-Cre::DYRK1A^flox/flox^ mice in which cardiomyocyte-specific deletion of DYRK1A starting in development had baseline cardiac hyperplasia and an increase in the expression of cell cycle genes (*11*). To complement LCTB-92 inhibitor experiments, we investigated the effects of the post-developmental, cardiomyocyte-specific ablation on DYRK1A in adult mice that underwent I/R MI. cTnnt2-Cre^Ert2^/+::Fucci2aR/Fucci2aR:DYRK1A^flox/flox^ (DYRK1A K/O) or cTnnt2-Cre^Ert2^/+::Fucci2aR/Fucci2aR (Control) mice were pulsed with tamoxifen for 10 days followed by a 2-week recovery period and then underwent an I/R MI with serial echocardiograms (**Fig. 7A**). DYRK1A K/O had improvements in LVEF at 2- and 4-weeks after MI compared to controls (**Fig. 7B & C**). Similar to LCTB-92 treated mice that underwent I/R MI, there were no significant differences in infarct size between our DYRK1A K/O and control groups (**Fig. 7D & E**). We have previously shown that bulk RNA-sequencing of isolated CMs from 7 days post-MI aMHC-Cre::DYRK1A^flox/flox^ mice demonstrated upregulation of mitotic and cell cycle-associated genes (*11*). Although methodological constraints prevented testing whether improvement in ventricular function from post-developmental DYRK1A knockdown is dependent on cell cycling (due to our knockout and αDKRC::DTA systems relying on CRE recombinase); we can infer that it most likely follows this mechanism based on our LCTB-92 DTA experiments. Together, the genetic ablation of DYRK1A in adult CMs recapitulated the effects of LCTB-92 treatment after MI, further supporting the potential therapeutic benefits DYRK1A inhibition after MI.

**Figure 7:**
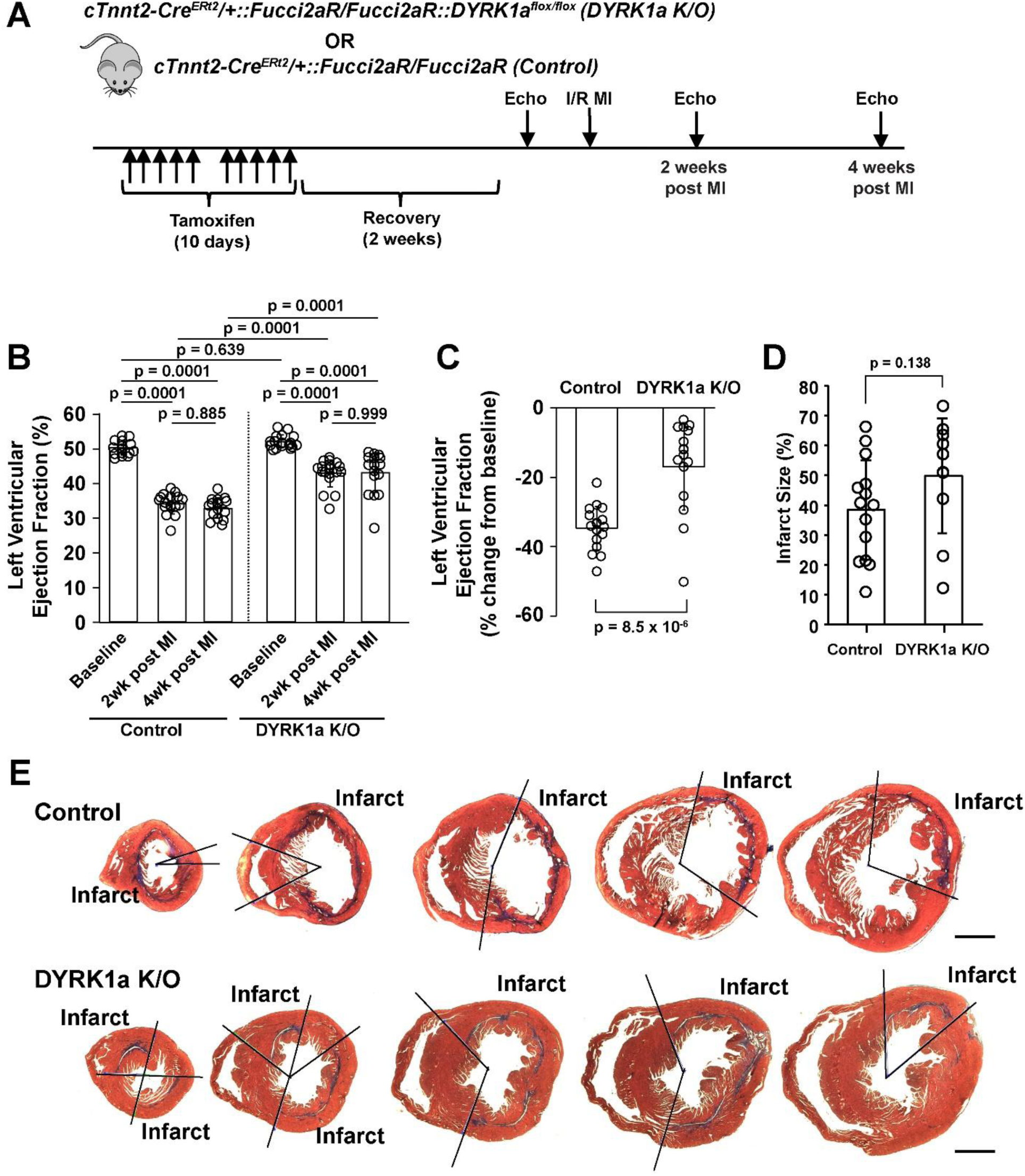
Post-developmental cardiomyocyte-specific deletion of DYRK1A recapitulates improved heart function induced by LCTB-92 treatment after I/R MI. **(A)** Schematic of 12-week-old cTnnt2-Cre^Ert2^/+::Fucci2aR/Fucci2aR::DYRK1A^flox/flox^ (Cardiomyocyte-specific DYRK1A K/O) or cTnnt2-Cre^Ert2^/+::Fucci2aR/Fucci2aR (Control) mice that undergo I/R MI. **B)** absolute (%), and **(C)** percent change (% change from baseline) LVEF 2- and 4-weeks post-MI. p-values were calculated using two-way ANOVA followed by Bonferroni’s multiple comparisons test. N = 18 DYRK1a K/O animals (11 males and 7 females) and N = 17 Control animals (9 males and 8 females). **(D)** Average infarct size (%) across whole heart sections from DYRK1A K/O or control hearts 4 weeks after MI with **(E)** representative images. N = 12 (6 males and 6 females) DYRK1A K/O animals and N = 14 (7 males and 7 females) Control animals. p-value by two samples t-test.

## DISCUSSION

Here, we mechanistically propose how small molecule inhibition of DYRK1A leads to increased cell cycling of cardiomyocytes both *in vitro* and *in vivo*. Our network model accurately predicts DYRK1A-mediated cell cycle activity that we leveraged to inform our *in vitro* validation experiments in NRCMs. Treatment of NRCMs with LCTB-21 and LCTB-92 exhibited a dose-dependent response on both cyclin D1 and Ki67 expression validating our model predictions. RNA-sequencing of LCTB-21 and LCTB-92 treated NRCMs revealed distinct transcriptional regulation of cell cycling genes, which we mapped to our network model’s predicted node activity via E2F1. LCTB-92 treatment induced CM cycling and improved heart function after I/R MI but did not alter infarct size. However, the beneficial effects of LCTB-92 treatment on heart function were lost when we genetically ablated cycling CMs, indicating that the functional benefits are mediated through cycling CMs. The post-developmental genetic ablation of DYRK1A attenuated the left ventricular dysfunction observed after MI, similar to the effects of LCTB-92. The two complementary approaches to prevent DYRK1A activity by pharmacological inhibition or genetic ablation specifically in adult CMs improved cardiac function after MI, supporting that DYRK1A knockdown after injury can lead to therapeutic benefits.

Our network model was built using literature in various cell types, as DYRK1A’s regulation of cell cycling is believed to be highly conserved. The model predicted experimental outcomes reported in cardiomyocyte literature not used to develop the model, despite the use of predominately non-cardiomyocyte literature to construct the model. Interestingly, while the model retained high accuracy against experiments from most cancer literature, the model failed to recapitulate experimental results when knocking down or overexpressing DYRK1A (*31, 38*). The inconsistency of the model to predict DYRK1A perturbations in certain cancer cells could be due to the type of cancer cell studied, where DYRK1A has been reported to be an oncogene. For example, DYRK1A knockdown reduced tumor growth in both *in vitro* and *in vivo* models of colon, breast, and neck cancers (colon, breast, neck) (*49*).

We show here that while LCTB-92 promotes CM cycling NRCMs and after I/R MI *in vivo*, it does not induce cycling in uninjured adult hearts. This could be due to the hypoxic environment created after injury. Several reports have demonstrated hypoxic environments can induce CM proliferation. Puente et al. (2014) showed that by exposing NRCMs to hypoxia, they could extend their proliferative window by reducing oxidative stress and DNA damage signaling (*50*). Similarly, Johnson et al. (2023) showed that chronic systemic hypoxia in adult mice induced limited CM proliferation in the right ventricle (*51*). These findings suggest that the post-injury hypoxic microenvironment may sensitize CMs to DYRK1A inhibition, thereby enhancing the proliferative response observed with LCTB-92 after MI.

The gene dosage of DYRK1A has also been observed to play an important role in CM cycling. Lana-Eloa et al. (2024) showed that Dp1Tyb mice, a line that has three copies of chromosome 21, including DYRK1A, and phenocopies human Down Syndrome, have proliferative defects in the embryonic heart (*14*). Flow cytometry showed an increase in the percentage of CMs in G1 phase, indicating a larger quiescent cell population. Additionally, they observed reduced p-RB1 and transcription of E2F target genes, consistent with our model predictions. Lastly, RNA-sequencing showed that LCTB-21 was able to partially rescue the decreased expression of proliferative pathways in Dp1Tyb embryonic hearts. Treatment with LCTB-92 might be able to induce a more robust recovery of these defects.

The αDKRC mouse model restricts Cre expression to adult CMs that express Ki67. One limitation is that the reporter mouse cannot distinguish between CMs that enter the cell cycle and undergo endoreplication to increase ploidy and CMs that undergo complete mitosis and cytokinesis to promote new daughter cells. Our prior investigations demonstrated that ∼10% of adult CMs that enter the cell cycle undergo bona fide proliferation as determined by pairs of clusters of Ki67-expressing CMs (*48*). With this in mind, we have several hypotheses of how the induction of CM cycling is responsible for the improvement of heart function we see with the treatment of LCTB-92 after I/R. LCTB-92 could be inducing complete CM proliferation that can restore some of the lost myocardium induced by I/R injury and improve the contractile function of the heart. Alternatively, LCTB-92 could also result in the induction of more polyploid CMs which have been shown to be more responsive to stress through enhanced mitophagy (*52*). Lastly, cycling CMs may produce paracrine factors that are cardioprotective to the surrounding myocardium. While the use of our αDKRC::DTA mice provides evidence that LCTB-92 improves ventricular function in a CM cell cycling dependent manner, we are unable to truly parse out what aspects of the cycling are beneficial for the myocardium. Additionally, while LCTB-92 administration may improve ventricular function through a combinatorial effect on multiple cell types, our αDKRC::DTA mouse experiments demonstrate that DYRK1A inhibition must act at least in part within CMs to confer functional benefit. Further work is necessary to understand the contributing role of LCTB-92 induced cycling CMs in recovery following I/R MI.

The use of our αDKRC::DTA mouse to show LCTB-92’s effect on ventricular function is dependent on CM cell cycling highlights the sensitivity of this subpopulation of CMs. Pride et al. (2010) demonstrated in a retrospective analysis of MI patients that there was no significant change in left ventricular function in individuals with infarct sizes <15% (*53*). These findings suggest that a substantial loss of CMs can occur without a measurable decline in ejection fraction. In our study, a marginal reduction of just 0.14% of the total CM population attenuated the benefit of LCTB-92 treatment, highlighting the functional importance and sensitivity of cycling CMs. Cycling CMs, as identified in the αDKRC system, are enriched for cells that have engaged coordinated cell-cycle and stress-response programs in vivo, a state that is rare in the adult heart and strongly associated with injury adaptation. These cells are therefore more likely to participate in repair-associated processes, including cytoskeletal remodeling, paracrine signaling, and maintenance of tissue integrity under stress.

It has also been reported that DYRK1A inhibitors trigger the proliferation of pancreatic B-cells both in isolated islet cells and in animal models of type 2 diabetes (T2D) (*54–56*). In addition to DYRK1A inhibition, DYRK1B inhibition appears to contribute to the beneficial effects of the inhibitors used in the diabetic contexts (*55*). Several structurally different DYRK1A inhibitors have been used including harmine (*57, 58*), and LCTB-92. *In vivo* treatment of the Goto-Kakisaki T2D rat model leads to proliferation of pancreatic β-cells and improvement of glycemic control. Like the injury-dependent effect of adult CM cycling we see here, the *in vivo* effects of LCTB-92 on β-cell proliferation are only observed in T2D pathological conditions and not in healthy controls. In both our animal model and the T2D animal models, it appears that LCTB-92 acts to restore homeostasis under pathological conditions.

Most DYRK1A inhibitors also inhibit DYRK1B, as well as some closely related kinases such as the cdc2-like kinases CLK1, CLK4 and, for some of them, more distantly related kinases like GSK3(*59*). LCTB-92 inhibits DYRK1A (IC50 value of 1.2nM), DYRK1B (1.8 nM), DYRK2 (40.2 nM), DYRK3 (19.3 nM), DYRK4(117.4 nM) as well as CLK1 (9.2 nM), CLK2 (0.6 nM), CLK3 (811.6 nM), CLK4 (6.0 nM), and GSK3β (4,158 nM)(*60*). Inhibition of some of these kinases may thus contribute to the effects of the LCTB-92 on CM cycling and heart remodeling after MI. The contribution of DYRK1B inhibition is rather likely as shown by Bayram et al. (*61*), and Zhang et al (*62*). Furthermore, CLKs are known to regulate alternative splicing, the modulation of which may affect CM cycling (*63*). The effects of genetic knockdown of DYRK1A on recovery after MI suggests the effects of LCTB-92 are mostly mediated through pharmacological inhibition of DYRK1A. Yet, inhibition of a few other kinases may complement DYRK1A inhibition in triggering the observed beneficial effects seen following MI.

The potential of the LCTB class of DYRK1A inhibitors to treat acute MI in patients is very attractive. Currently, LCTB-21 is being investigated in a first-in-human Phase 1 trial in healthy volunteers and subjects Down Syndrome and Alzheimer’s disease (Clinical Trial NCT06206824). While we show that both LCTB-21 and LCTB-92 induce cycling in CMs *in* vitro, LCTB-92 showed stronger cell cycling effects which led to our investigation of LCTB-92 *in vivo*. Whether LCTB-21 has similar effects as LCTB-92 on cardiac function and CM cycling after MI remains to be seen. Importantly, we did not observe tumorigenic effects of LCTB-92 in our studies. The results of this trial, in conjunction with our results using LCTB-92 in a preclinical animal model of MI, are encouraging in that the inhibition of DYRK1A may represent a new first-in-class therapy to treat heart disease. Follow up studies that recapitulate the results here in more human relevant model systems would be necessary to investigate the potential of these compounds in treating MI.

## MATERIALS AND METHODS

### Study Design

The aim of this study was to evaluate if a specific oral inhibitor of DYRK1a would improve left ventricular function after myocardial infarction and increase adult cardiomyocyte cycling. The study used cultured neonatal rat cardiomyocytes (NRCMs) to examine Ki67 expression in response to LCTB-92, LCTB-21, and their inactive isomers. RNAseq was used to investigate changes in the gene expression of NRCMs in response to LCTB-92 and Iso-LCTB-92. For in vivo studies, we used male and female to investigate sex as a biological variable. We used *αDKRC::RLTG* mice that restrict Cre expression to adult mouse cardiomyocytes that re-enter the cell cycle. *αDKRC::DTA* mice were used to express Diphtheria toxin and ablate in adult cardiomyocytes that re-enter the cell cycle. *cTnnt2-CreERt2::DYRK1a^flox/flox^* mice were used to investigate the genetic ablation of DYRK1a in adult cardiomyocytes. Ischemia-reperfusion myocardial infarctions were performed by ligating the proximal left anterior descending coronary artery for 60 minutes, followed by restoration for blood flow. This model recapitulated acute myocardial infarction and reperfusion in humans. Cycling cardiomyocytes were quantified by counting eGFP+ cells in 10-micron short-axis histological heart sections from *αDKRC::RLTG* mice. Infarct sizes were quantified by examining serial sections stained with Masson trichrome. Sample sizes were selected based on previous experience with similar methods. Mice were randomized to treatment groups based on genotypes for each experiment. Investigators were blinded to genotype when performing surgeries, echocardiography, quantification of eGFP+ cardiomyocytes, and analyses of infarct sizes. No samples were excluded from the study. Details on sample sizes representing biological replicates and statistical tests are detailed in figure legends and the Statistical Analysis section of the Materials and Methods.

### Network Model Construction

A logic-based differential ordinary equations network model was used to predict mechanisms of DYRK1A-mediated cell cycle activity. The network model was built using Netflux (*23*), a software that auto-generates a logic-based differential equation model based on a list of reaction rules and parameters (*22*). This includes assigning the interactions (inhibitory or stimulatory) between molecular components of interest (nodes) as well as the logic-gating strategy (“OR” or “AND” gates). Further information on how this modeling approach works can be found in Kraeutler et al. (2010) and Clark et al. (2024). A series of 6 primary and review literature articles (**Table S1**) were used to infer the interactions and logic-gating strategy within the model (**Table S2**). Literature was chosen based on its relevance to DYRK1A-mediated cell cycle control and dimerization partner, RB-like, E2F and multi-vulval class B (DREAM) complex regulation. While literature that used cardiomyocytes or other relevant experimental models were preferred, we used literature across multiple experimental models (noted in **Table S1**) as this regulatory process is believed to be highly conserved across tissue types. Literature search key terms included: proliferation, cell cycling, DREAM complex, and DYRK1A.

Default reaction parameters included reaction weight (w=1), Hill coefficient (n=1.4) and half-maximal effective concentration (EC_50_=0.50) and default node parameters included initial activation (Y_init_=0), maximal activation (Y_max_=1), and time constant (τ =1). Parameter optimization was run on input nodes to allow the model to experience change with perturbations at baseline. DYRK1A and B-Myb were then set to a Y_max_=0.9 and 0.1 respectively to establish baseline activity. As in past logic-based differential equation network models (Kraeutler et al., 2010; Tan et al., 2017; Zeigler et al., 2016) species (nodes) refer to a small molecule, gene, protein, or process. Reactions (or edges) are activating and inhibiting relationships between network species (*22, 64, 65*).

### Network Model Validation

Model simulations were compared to experimental data from primary literature that was separate from the literature that was used to build the model’s interactions and logic-gating strategy. Guided by the experimental design of each validation study, we simulated knockdowns or overexpression of specific nodes to see how these perturbations could affect the DNA replication node within the model. Knockdowns were performed by setting the Y_max_ of a desired node to 0. Overexpression was performed by setting Y_max_= 10 for input nodes or by setting Y_init_= 10 and tau= 10^9^ for intermediate nodes. All perturbations were performed after the model reached a steady state (<0.05% change in activity for all nodes). Model accuracy was determined by if simulation results qualitatively matched the statistics from the corresponding experiment in the primary literature (n = 19). Validation was performed across cancer and cardiomyocytes to look at deviations in model predictions across different biological contexts. Experimental figures were replotted with WebPlotDigitizer (*66*).

### Network Model Simulations

We simulated a concentration response by knocking down DYRK1A in increments of 1%. We then looked at the change in activity of the responding nodes to characterize their sensitivity to a DYRK1A knockdown. EC50 and IC50 of affected nodes were determined by the % DYRK1A knockdown at which their normalized activity was changed by 50%. A knockdown screen was conducted through simulated 100% knockdowns on each node of the model at baseline activity, where DYRK1A is highly active (Y_max_=1). We then examined the change in activity of the network in response to each individual knocked down node. We simulated “low, medium, and high” DYRK1A inhibitor concentrations by knocking down DYRK1A in increments of 15% to simulate “low, medium, and high” drug concentrations. Our concentrations corresponded to these simulated knockdown values; low: 15%, medium: 30%, and high: 45%. The results were then compared to in vitro data of neonatal rat cardiomyocytes that received the LCTB-92 and LCTB-21 DYRK1A inhibitors.

### Neonatal rat cardiomyocyte isolation, immunostaining, and image analysis

Cardiomyocytes were isolated from 1-2-day-old Sprague-Dawley rats using a Neomyt isolation kit (Cellutron). Each cell isolation is a mixture of cardiomyocytes from all 9-12 male and female pups from that litter. The cells were cultured in plating media (low-glucose Dulbecco’s modified eagle media (DMEM), 17% M199, 10% horse serum, 5% fetal bovine serum (FBS), 1% L-Glutamine, 10 U/mL penicillin, and 50 mg/mL streptomycin) at a density of 30,000 cells per well in a 96-well Corning CellBIND plate coated with SureCoat (Cellutron). After being cultured for 48 hours, the cells were serum-starved in WE+B medium (William’s E medium without phenol red (Gibco) supplemented with Primary Hepatocyte Cell Maintenance Cocktail B (Gibco) containing P/S, ITS+, GlutaMAX, and HEPES) for 4 hours before drug treatment. Cardiomyocytes were then treated with the DYRK1A inhibitors LCTB92 or LCTB21 (0.01-1 µM) or respective isomeric and DMSO (0.01%) controls in serum-free media for 48 hours.

Cells were then fixed in 4% paraformaldehyde and fluorescently labeled with anti α-actinin (Sigma) or cardiac troponin T (Abcam), anti-Ki67 (Invitrogen), anti-cyclin D1 (Abcam), and DAPI (Life Technologies). Stained cells were imaged with the Operetta CLS high-content imaging system using a 10x 0.3 NA objective. Image analysis was performed using a CellProfiler pipeline for cell segmentation to quantify the percentage of cardiomyocytes positive for Ki67 or cyclin D1 (*67*). Thresholds for mean nuclear Ki67 or cyclin D1 intensity were set manually to approximate the local minimum of the bimodal distribution. Multinucleated cardiomyocytes were identified by measuring the distance between adjacent nuclei, with a threshold of less than 2 pixels. Cell cycle phase was assessed by combining DNA content analysis and Ki67 positivity as described in previous studies (*68, 69*).

### Gene Expression Analysis

NRCMs were isolated and cultured as previously described, then seeded at a density of 1,000,000 cells per well in 6-well Corning CellBIND plates coated with SureCoat (Cellutron). Cells were treated with 1 µM of either LCTB-92 or LCTB-21, their respective isomeric controls, or DMSO (n = 3 per condition), following the same treatment protocol as described above. RNA extraction, cDNA library preparation (including adapter ligation), and sequencing were performed by Genewiz, Inc. using an Illumina® HiSeq® system. FASTQ files were processed using a Nextflow pipeline (STAR + Salmon), in which reads were aligned with STAR and transcript abundance was quantified using Salmon (*70*). Principal component analysis (PCA) and differential gene expression were conducted using the **DESeq2** package in R (*71*). Gene Ontology (GO) enrichment analysis was performed with the **ClusterProfiler** package in R (*72*). Transcription factor enrichment was assessed using the **ChEA3** API platform, with transcription factors ranked by average rank across all libraries (*45*). To integrate RNA-seq data with our network model, we linked the E2F1 node to its target genes identified via ChEA3 and mapped these to gene sets associated with DNA replication, cytokinesis, and cardiac contraction GO terms.

### Mice

*αDKRC* (*<u>α</u>MHC-Mer<u>D</u>reMer-<u>K</u>i67p-<u>R</u>oxed<u>C</u>re::<u>R</u>ox-<u>L</u>ox-td<u>T</u>omato-e<u>G</u>FP) mice were made and maintained as previously described. C57BL/6J* (JAX stock #000664), *RC::RLTG* (*B6.Cg-Gt(ROSA)26Sor^tm1.2(CAG-tdTomato,-EGFP)Pjen^/J*), and *ROSA-DTA* (*B6.129P2-Gt(ROSA)26Sor^tm1(DTA)Lky^/J*) mice were purchased from Jackson labs. *CAG-STOP-Fucci2aR* was obtained from the European Mouse Mutant Archive. *cTnnt2-Cre^ERt2^* was provided by Dr. Chenlen Cai. The *αDKRC::RLTG*, *αDKRC::DTA*, *cTnnt2-Cre^ERt2^::Fucci2aR*, and *cTnnt2-Cre^ERt2^::Fucci2aR::DYRK1a^flox/flox^* mice were created by standard breeding methods and all mice were genotyped. DYRK1A knockdown was confirmed by qPCR in our previous manuscript (*11*). We were unable to show protein level knockdown of DYRK1A due to the known limitations of DYRK1A antibody performance in tissue lysates.

### Compounds

LCTB-92, ISO-92, LCTB-21, and ISO-21 were provided by Perha Pharmaceuticals. For experiments using NRCMs, LCTB-92, ISO-92, LCTB-21, and ISO-21 were dissolved in DMSO and added to cell media to give a final contraction of 0.01 μM, 0.1 μM, or 1 µM. For in vivo experiments, LCTB-92 or ISO-92 was dissolved in 0.5% carboxymethylcellulose in sterile water and 1 mg/kg was given daily by oral gavage.

### Left Anterior Descending (LAD) I/R MI surgery

All animal care and surgeries were in accordance with UVA ACUC Policy on Rodent Surgery and Perioperative Care under ACUC-approved animal protocol (UVA ACUC Wolf Lab protocol #4080). The individual performing surgeries was blinded to the mouse genotypes and treatments. Procedure: Male and female ten-to twelve-week-old mice were randomized to MI surgeries, weighed and then anesthetized in an induction chamber using the gas anesthetic Isoflurane (3% volume/weight, oxygen 500ml/min). The animal was placed in the supine position on a face mask connected to the anesthesia system. Isoflurane was adjusted to provide a maintenance level (1.8-2.2% volume/weight, oxygen 500ml/min) throughout the procedure. Anesthesia was monitored closely by the following methods: (1) Depth and rate of respiration; (2) Heart rate by ECG; (3) Mucous membrane color; (4) Body Temperature via Physio Suite Homoeothermic Temperature Monitoring System; (5) Reflexes & toe pinch; (6) Overall appearance of muscle relaxation. All surgical procedures were carried out with a stereo microscope exclusively for small animal surgery. Normal body temperature were maintained using an electrical heating pad on a feedback system via rectal probe.

To prepare for surgery, the mouse was first given subcutaneous fluids (veterinary Normosol), and Atropine as a pre-anesthetic to decrease mucous secretion and to prevent gag reflex during intubation. The neck and chest wall were shaved, and then prepped three times with alternating wipes of povidone-iodine and 70% ethanol. Mice were placed in the supine position on a heating pad and an endotracheal intubation was performed under direct laryngoscopy with a lighted fiber optic stylus to visualize the vocal cords and insert tracheal tube. The endotracheal tube was then connected to the automated ventilator (Kent Scientific, Inc.) and the mouse was ventilated (tidal volume = 1.0 mL, rate = 120 breaths/minute). Bupivacaine was infiltrated subcutaneously into the area of the left 3rd and 4th intercostal space. A left anterior thoracotomy was performed using sterile technique by making a small subcutaneous incision lateral from the sternum with sharp scissors. The pectoralis muscle groups were identified and separated by blunt dissection, and held open using fixed retractors. The thoracic cage was exposed and the 3rd intercostal space was identified. Incision into the 3rd intercostal space was done by a combination of blunt micro-scissors and micro-cauterizing tool in order to minimize bleeding. Retractors were then repositioned onto the upper and lower ribs in order to visualize the heart. The pericardium was then blunt dissected in order to expose the anterior view of the heart. The LAD on the surface of the left ventricular wall was identified and an 8-0 prolene suture was placed through the myocardium into the anterolateral LV wall underneath the LAD at the level of the lower atrium (1 mm below the left auricle), and the suture was tied. Occlusion of blood flow is observed by blanching of the heart muscle and ST elevation confirmed by ECG. After 60 minutes the ligature was removed. The chest wall, facial planes, and skin were sutured and the mouse was recovered and treated with analgesics. In sham control mice, the entire procedure was identical except for the ligation of the LAD. The mortality rate associated with surgeries was ∼10%.

### Histology and Immunohistochemistry

Hearts were excised and fixed in 10% Neutral Buffered Formalin (Fisher, Inc.) for a minimum of four hours prior to embedding in paraffin. Ten-micron sections were prepared in short axis orientation by microtome with 8 sections per glass slide. Paraffin was removed and the tissue sections were rehydrated using Xylene and serial ethanol wash steps, respectively. Antigen retrieval was performed by incubating tissue sections in boiling 1x Unmasking solution (Vector Labs H-3300) for 22 minutes. After cooling to room temperature, the tissue sections were treated with Sudan Black to quench auto-fluorescence. Briefly, tissue sections were incubated in 0.1% Sudan Black (Sigma, Cat# 199664) in 1x PBS and 70% ethanol at room temperature for 20 minutes to quench auto fluorescence, followed by 3 x five-minute washes in 1x PBS containing 0.02% Tween 20, and a final five-minute wash in 1x PBS.

### Quantification of cardiomyocyte cycling

Images were obtained using a Leica DM2500 Fluorescence microscopy system with a Leica DFC7000 T fluorescence color camera or a Leica THUNDER imaging system and Leica LAS X Multi Channel Acquisition software. For imaging, fluorescence channels were calibrated to background of control sections stained with secondary antibodies alone. eGFP+ CMs visualized using an YFP filter and quantified from 6-8 ten-micron short axis sections of three slides per animal separated by ∼400 microns per slide. The infarct zone was identified by WGA. The non-infarct zone was divided into thirds with the two regions adjacent to the infarct defined as the border zones and the middle region defined as the remote zone. eGFP+ CMs present in the same location on sequential slides were counted once among all sections to avoid over-representation of eGFP+ cells.

### Echocardiography

Mice were anesthetized with Isoflurane and gently restrained on a Vevo integrated rail system. The system included a physiological monitoring unit consisting of a heating board with integrated ECG electrodes. The table temperature was maintained at ∼38°C using a rectal probe for temperature feedback. Hair was gently removed, and ultrasound contact gel (warmed to 37°C) was applied to the chest. Echocardiography was performed using a high frequency 30 MHz linear transducer and a Vevo 1100 (Visual Sonics) system similar to previously described methods(*47, 48*). Initial B-mode images were obtained in the parasternal long axis with the apex and aortic valves visualized. Next, B-mode short axis images were obtained by turning the ultrasound probe ∼90 degrees and identifying the mitral valve papillary muscles. The parasternal long axis images were analyzed using VevoView software to calculate left ventricular dimensions. The individuals performing and analyzing the echocardiography were blinded to the treatment groups.

### Statistics

GraphPad Prism 9 (GraphPad Software, Inc.) was used for statistical analyses. The data was analyzed for normal distribution using Anderson-Darling, D’Agostino & Pearson, Shapiro-Wilk, and Kolmogorov-Smirnov tests in GraphPad. One-way ANOVAs with Tukey test corrections for multiple comparisons, two-way repeated measured ANOVA with Bonferroni test for multiple comparisons, and student t-tests were used. We used a calculator from the IACUC at Boston University, including an Excel template for calculation based on means/standard deviations and proportions (https://www.bu.edu/researchsupport/compliance/animal-care/working-with-animals/research/sample-size-calculations-iacuc/). For experiments quantifying eGFP+ CMs, we assumed a Type I (alpha) error of 0.05, a difference (delta) of 2-fold (∼2-3 eGFP+ cells in sham heart sections and 4-6 eGFP+ cells in MI heart sections), a standard deviation of 2-3, and a Power of 0.9. We calculated that nine mice would be needed in each group. For the echocardiography experiments, we assumed a Type I (alpha) error of 0.05, a difference (delta) of ∼10% (EF ∼30% in the MI control group and ∼40% in the drug treated group), a standard deviation of 5%, and a Power of 0.95. We calculated that a minimum of ∼seven mice would be needed in each group. We found no statistically significant evidence for sex differences in any of our *in vivo* experiments.

### Study Approval

All mice were housed and maintained in accordance with UVA Animal Care and Use Committee approved protocols (UVA Animal Care and Use Committee (ACUC) Wolf Lab protocol #4080). Rats were housed and maintained in accordance with UVA Animal Care and Use Committee approved protocols (UVA Animal Care and Use Committee (ACUC) Saucerman Lab protocol #4080).

## Data availability

All data associated with this study are present in the paper or Supplementary Materials. RNAseq data will be deposited into the Gene Expression Omnibus (GEO) maintained by NCBI. Netflux code is available at https://github.com/saucermanlab/Netflux. Mouse data is available by request.

## Author contributions

J.J.S. and M.J.W. designed the study, obtained funding, supervised the work, and edited the manuscript. B.C.M. performed the network modeling, RNA sequencing, and experiments on tissue sections, and wrote the manuscript. K.L.W. performed the in vitro experiments, image analysis, and wrote those sections of the manuscript. B.N.H., M.W., and C.Z. performed network modeling. A.Y, L.A.B., K.S., D.H., L.M., and M.J.W. designed, performed, and analyzed the mouse surgery data, echocardiography, histology, and immunohistochemistry. A.Y. performed the mouse surgeries and echocardiograms. L.M., M.F.L., and E.D. provided the synthesized and purified LCTB-92, ISO-92, LCTB-21, and ISO-21 and expertise regarding the design of the mouse experiments.

---

**Figure S1:**
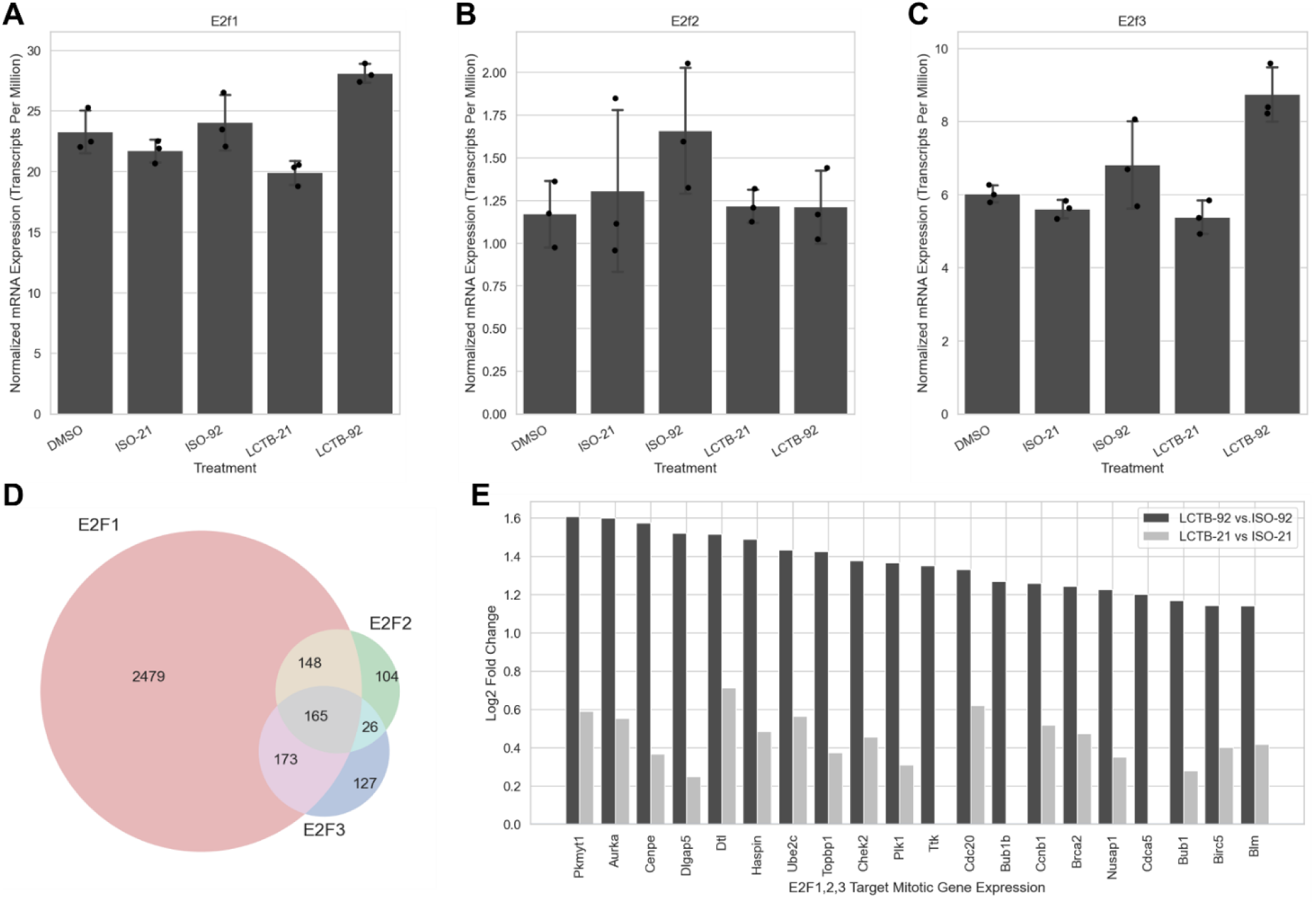
Increased E2F1/2/3 activity in drug-treated NRCMs. (A–C) Normalized mRNA expression of **(A)** E2F1**, (B)** E2F2, and **(C)** E2F3 across treatment groups in neonatal rat cardiomyocytes (NRCMs). **(D)** Venn diagram of predicted E2F target genes identified using ChEA3 (mean rank library). **(E)** Significant differentially expressed mitotic genes (as determined by GO term enrichment gene sets) that are predicted E2F targets in LCTB-92 or LCTB-21-treated NRCMs relative to respective isomer controls. Statistical significance for panels **A–C** was assessed using one-way ANOVA followed by post hoc pairwise comparisons to DMSO with Benjamini-Hochberg correction for multiple testing.

